# A synthetic skin bacterial community (SkinCom) enables reproducible investigations of the human skin microbiome

**DOI:** 10.1101/2024.06.01.596969

**Authors:** Asama Lekbua, Deepan Thiruppathy, Joanna Coker, Yuhan Weng, Fatemeh Askarian, Armin Kousha, Clarisse Marotz, Amber Hauw, Victor Nizet, Karsten Zengler

**Affiliations:** Division of Host-Microbe Systems & Therapeutics,Department of Pediatrics, University of California, San Diego, 9500 Gilman Drive, La Jolla, CA 92093-0760, USA; School of Biological Sciences, University of California, San Diego, La Jolla, CA, 92093-0376, USA; Department of Bioengineering, University of California, San Diego, La Jolla CA 92093-0412, USA; Bioinformatics and Systems Biology Program, University of California San Diego, La Jolla, California, 92093-0419, USA; Glycobiology Research and Training Center, UCSD, La Jolla, CA, USA; Skaggs School of Pharmacy and Pharmaceutical Sciences, UC San Diego, La Jolla, CA, USA; Center for Microbiome Innovation, University of California, San Diego, La Jolla, CA 92093-0403, USA; Program in Materials Science and Engineering, University of California, San Diego, 9500 Gilman Drive, La Jolla, CA 92093-0418, USA

**Keywords:** Skin microbiome, synthetic community (SynCom), host-microbe, cosmetic compounds, sodium lauryl sulfate, sodium lauryl ether sulfate, rhamnolipids, creatine, epicutaneous murine model

## Abstract

Existing models of the human skin have aided our understanding of skin health and disease. However, they currently lack a microbial component, despite microbes’ demonstrated connections to various skin diseases. Here we present a robust, standardized model of the skin bacterial community (SkinCom) to support complex *in vitro* and *in vivo* investigations. Our methods lead to an accurate, reproducible, and diverse community formation of aerobic and anaerobic bacteria. Subsequent testing of SkinCom on the dorsal skin of mice allowed for DNA and RNA recovery from both the applied SkinCom and from dorsal skin, highlighting its practicality for *in vivo* studies and -omics analyses. Furthermore, our results indicate that 65.6% of the responses to common cosmetic chemicals *in vitro* were recapitulated in a human trial. Therefore, SkinCom represents a valuable, standardized tool for investigating microbe-metabolite interactions and facilitates the experimental design of *in vivo* studies targeting host-microbe relationships.

## Introduction

Human skin, the body’s largest and most exposed organ, hosts a diverse microbiota critical for defense against microbial pathogens and other skin pathologies^1–5^. Disruptions to the skin microbiome can lead to diseases, such as acne vulgaris, atopic dermatitis, and seborrheic dermatitis. Furthermore, the application of topical products, including make-up^6^, deodorant^7^, and various skin care remedies can alter the microbiome’s composition and diversity for weeks^8^. Understanding the impacts of skin microbiome dysbiosis, such as through skincare products and their ensuing effects on skin health, support the development of personalized skin treatment regimens.

The composition of the skin microbial community is influenced by various microenvironmental factors, such as pH, temperature, moisture, oxygen availability, and topography^9–11^. Microbial interactions and intercellular metabolite exchanges play a crucial role in shaping the community’s structure and function^2,11,12^. Research on the human skin microbiome in disease states or in response to chemicals has improved our understanding of its role in skin health and facilitated targeted intervention studies^13–16^. However, the highly individualized nature of the skin microbiome, with variation in the distribution and gene expression profiles of identical microbes between individuals^10,17,18^, often complicates cross-study comparisons.

Establishing a model of the skin microbiome could enable reproducible studies of complex metabolite-microbe, microbe-microbe, and host-microbe interactions that govern skin health. However standardized studies of the skin microbiome are severely hampered by the lack of reproducible *in vitro* systems^19,20^. Despite dynamic modifications by external and internal factors, documentation of the skin microbiome composition across different body sites and time points reveals that its composition is largely represented by a select few resident microorganisms^10,13^. The recent advent of reconstructed human epidermis models capable of capturing certain key physiological complexities of skin, such as Epiderm, Labskin, or NativeSkin, has aided studies on microenvironmental regulation of microbial community structure^21–28^. Moreover, previous work by Uberoi et al. has constructed a synthetic skin community with 6 species of bacteria, from 3 genera^29^. However, it lacks the anaerobic *Cutibacterium* genus, which is abundant on the human skin^10,18,30^. Therefore, a robust standardized bacterial community for probing the community dynamics of the skin microbiome is still lacking.

In this study, we developed a nine-species bacterial synthetic community to represent the average human skin microbiome (SkinCom). This community was constructed using an automated, programmable microfluidic device employing piezo-electric technology for dispensing pico-liter droplets at nanoscale accuracy^31^. This method ensures that communities are defined by the number of cells at assembly time, yielding highly reproducible results. The approach supports high-throughput *in vitro* community generation, allowing for the study of many replicates quickly and cost-effectively. We assembled SkinComs from five starting inoculum proportions and used shotgun metagenomic sequencing to analyze community composition and diversity.

After establishing the reproducibility of SkinCom *in vitro*, we then assessed incorporation of the SkinCom into existing *in vivo* systems, specifically a murine epicutaneous model and human skin swab studies. In the murine epicutaneous model the SkinCom was applied onto the shaved dorsal skin of CD1 mice, with DNA and RNA recovered three days post-incubation and aspects of host skin barrier integrity investigated. In the human trial, we explored SkinCom’s growth, diversity, and composition change in the presence of cosmetic chemicals, comparing *in vitro* differential abundance results with those from a human trial involving skin applications of the same chemicals. This study thus establishes a model SkinCom suitable for both *in vitro* and *in vivo* investigations, demonstrating its utility in assessing the impact of chemical perturbations on the skin microbiome.

## Results

### Selection and growth characterization of SkinCom species

The composition of the skin microbiota varies significantly across body sites and among individuals^10,18,30^, yet it is typically dominated by a few microbial genera^10,13^. To model this diversity, we constructed a synthetic skin bacterial community (SkinCom) comprising eight bacterial species most frequently found across human skin^2^: *Corynebacterium afermentans* (ATCC 51403, strain CIP 103499 [LCDC 88199]), *Cutibacterium acnes* (ATCC KPA171202), *Micrococcus luteus* (ATCC 4698), *Staphylococcus capitis (ATCC 27840, strain LK 499), Staphylococcus epidermidis (*ATCC 12228)*, Staphylococcus hominis* (ATCC 27844, strain DM 122), *Staphylococcus warneri* (ATCC 27836, strain AW 25), and *Streptococcus mitis* (ATCC 49456, strain NCTC 12261). Given its profound significance in human health and disease, we also included *Staphylococcus aureus* (ATCC 35556, strain SA113)^14,15,32–35^. Each type strain was initially cultured in a complex nutrient-rich media (Brain Heart Infusion, BHI) at 37°C in the presence of oxygen, except for *C. acnes*, which required cysteine-reduced medium in hungate tubes due to its anaerobic nature. Upon reaching late log phase, cultures were diluted to an optical density (OD600) of 0.065 and arrayed into 96-well plates using a pico-liter printing microfluidic device. Plates were incubated at 37 °C in the presence of oxygen (oxic) or in the anaerobic chamber (anoxic) (Figure 1A).

**Figure 1.**
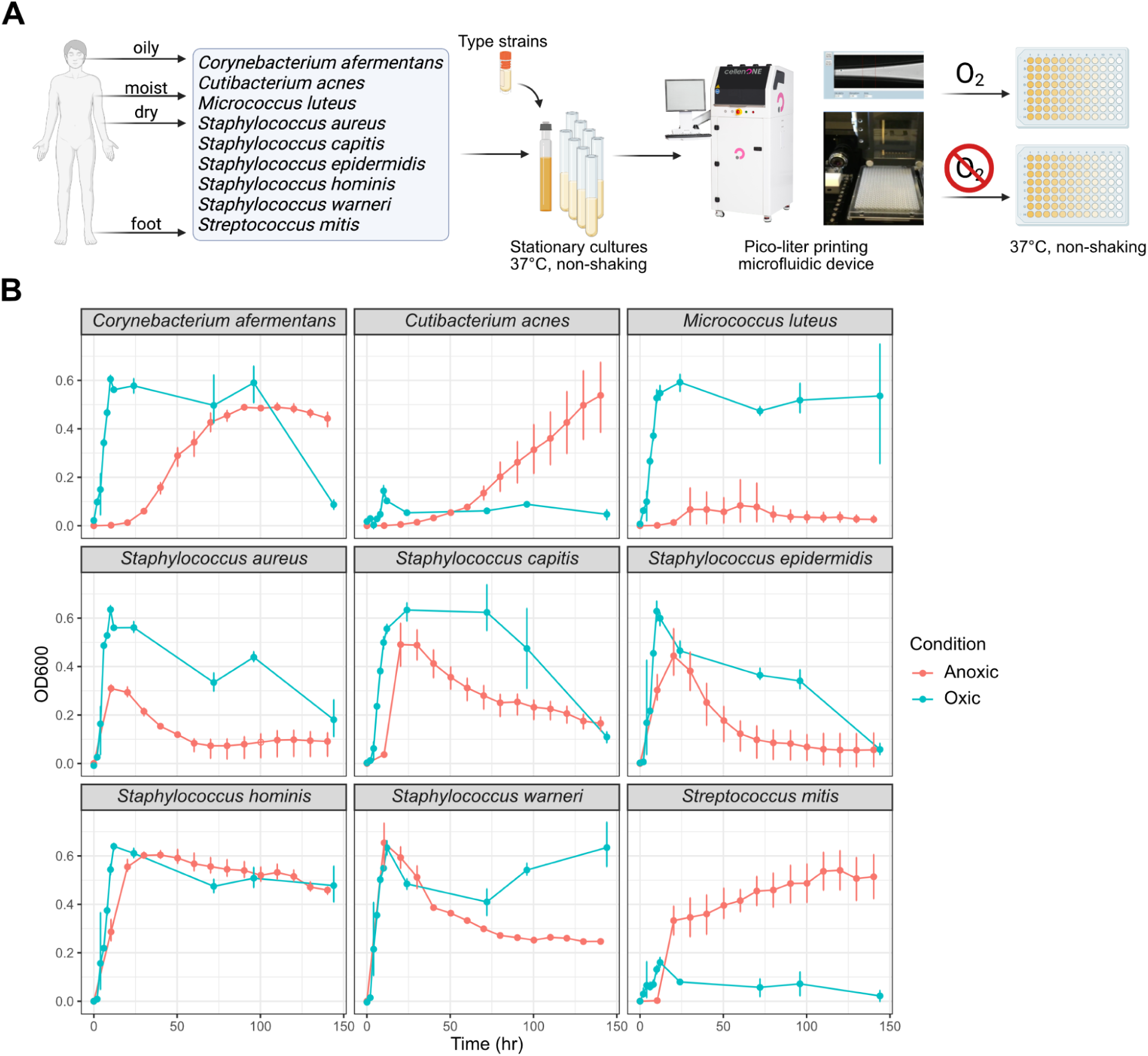
Experimental design and growth curves of SkinCom species. A) Bacterial species were selected to represent various microenvironments throughout the human body. Type strains were obtained from ATCC and cultured oxically, with the exception of *C. acnes* (anoxically). Cultures were then diluted to OD600 = 0.065 and arrayed into a 96-well plate using the pico-liter printing microfluidic device. One plate was incubated at 37 °C in the presence of oxygen, and another was incubated at 37 °C in the anaerobic chamber (see Methods). B) Growth of each species under aerobic and anaerobic conditions, measured by OD600 (n=4). Strains were grown in 200*μ*l of 1X BHI medium. Oxic and anoxic growth was monitored for 144 and 140 hrs respectively.

We monitored the growth rate of each species over 144 hours through optical density measurements. Although the surface of the skin is exposed to oxygen, the varying oxygen availability throughout the skin layers allows for anaerobic growth, exemplified by *C. acnes*^2,35^. We therefore tracked the growth rates of each SkinCom member under both oxic and anoxic conditions (Figure 1B). In oxic conditions, all species, except *C. acnes* and *S. mitis*, achieved a maximum OD600 above 0.2. Conversely, in anoxic conditions, all except *M. luteus* surpassed an OD600 of 0.3, with *S. warneri* showing the highest growth in both environments. The lowest growth under oxic and anoxic conditions was observed for *C. acnes* and *M. leuteus*, respectively. Growth rates were determined using the Growthcurver R package for each species^36^.

### Determining media conditions for SkinCom diversity

After assessing the growth rate of each species individually, we proceeded to combine them into a synthetic community. This step aimed to foster reproducible growth while maximizing the community’s diversity, ensuring no single species would dominate. To achieve this, each species was grown to OD600 = 0.065 then mixed with one another in a 1:1 ratio, forming an equally-mixed (EM) community. Using a Scienion CellenONE liquid printer, known for its precision in dispensing of 400 picoliters droplets, we added 200 droplets of each standardized culture to a 96-well plate containing 200 *μ*l of either 0.1x or 1.0x brain-heart infusion (BHI) media (n = 8). Consequently, each destination well comprises 200 *μ*l of media and 80 nL of the mixed bacterial isolates (200 drops of 400 picoliters). For negative controls, 80 nL of sterile BHI was added to separate wells (n = 8). The plate was incubated for 5 days at 37 °C oxically without shaking.

OD600 readings demonstrated consistent growth patterns within technical replicates, but revealed a distinction between communities grown in 1.0X and 0.1X BHI media (Figure 2A). Specifically, in the 1X BHI media, OD600 rose sharply within the initial 10 hrs, followed by a subsequent reduction and subsequent stabilization of growth levels between 48 to118 hrs. Conversely, communities in the 0.1X BHI medium exhibited a lesser peak OD600, maintaining steady levels from 24 to 48 hrs, before experiencing a decline thereafter.

**Figure 2.**
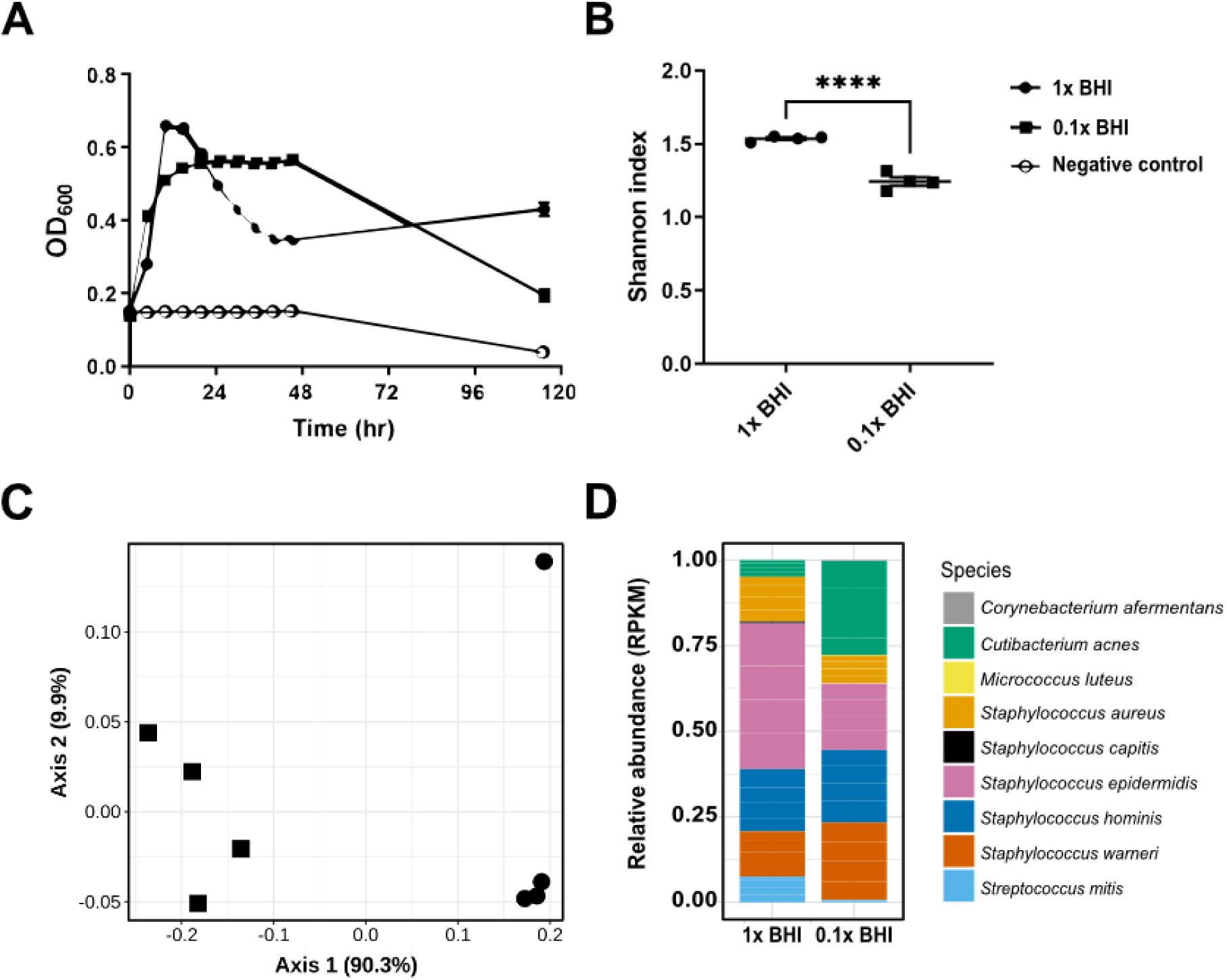
Determining the effect of the media condition for equal-mix SkinCom based on optical density and diversity. A) OD600 readings of communities grown in 1X or 0.1X BHI under anaerobic conditions (n = 8 per condition). Readings were taken every 5 hours. Data are represented as mean ± SEM. B) Shannon diversity index of 1X and 0.1X BHI communities at 120 hrs (n = 4 each), (Student’s t-test, p = <0.0001). C) PCA of Bray-Curtis distance of 1X and 0.1X BHI communities (n = 4 each), (PERMANOVA, p = 0.032). D) Relative abundance of 1X and 0.1X BHI communities (n = 4 each) at 118 hrs, determined by shotgun metagenomics sequencing (See Methods).

Following the assessment of growth rates and the establishment of the SkinCom, we proceeded to analyze its composition and diversity through shotgun metagenomic sequencing after 118 hours of growth (n = 4 per condition). This approach sought to quantify the alpha-diversity within our synthetic communities using the Shannon diversity index as a measure. Contrary to expectations based on prior studies with synthetic microbial communities from soil, where diluted media contributed to increased diversity^31^, we found that SkinComs grown in 0.1X BHI had significantly lower diversity than those grown in 1X BHI (Figure 2B). This discrepancy underscores a unique response of skin microbial communities to environmental conditions, differing from soil-based counterparts.

Further analysis employing the Bray-Curtis distance between samples highlighted the significant compositional differences between the 0.1X and 1X BHI communities (Figure 2C). Despite these variations in diversity and composition, sequencing data confirmed the presence of all intended isolates within both sets of SkinComs, indicating successful community assembly (Figure 2D). Strikingly, when *C. acnes* was cultivated axenically, no growth occurred as expected for this anaerobic bacterium (Figure 1B). However, when grown in the SkinCom, we were able to detect its presence. Importantly, metagenomic results confirmed the ability of *C. acnes* to proliferate in an oxic setting, likely facilitated by oxygen consumption by aerobic members of the SkinCom. Given these insights and the superior diversity observed in 1X BHI cultures, we decided to carry out all subsequent experiments under aerobic conditions using 1X BHI.

### Optimizing community diversity by altering the starting inoculum

Building on previous work with a rhizosphere synthetic community, which showed that alpha-diversity in the community could be enhanced by adjusting the starting ratios of individual microorganisms^31^, we explored this concept with the SkinCom. We generated community inocula with four distinct starting ratios, then compared the experimental setups to the standard equally-mixed (EM) SkinCom configuration. The design of these varied ratios generally intended to balance the community by incorporating fewer fast-growing and more slow-growing microorganisms, based on earlier growth curves of isolates (Figure 1). Specifically, we employed a 2x cutoff to categorize microorganisms as either fast or slow growers, and a 3x cutoff to further distinguish them into fast, moderate, and slow categories. Additionally, we adjusted the starting proportions based on the growth curve’s slope (GCS) and the time to reach the midpoint of growth (GCT) for each microbe, adding further precision to community assembly (Figure 3A)^36^. The absolute and relative proportions of the inocula are shown in Figure 3A and 3B. Details about the assembly of the community is provided in Supplemental Table S1.

**Figure 3.**
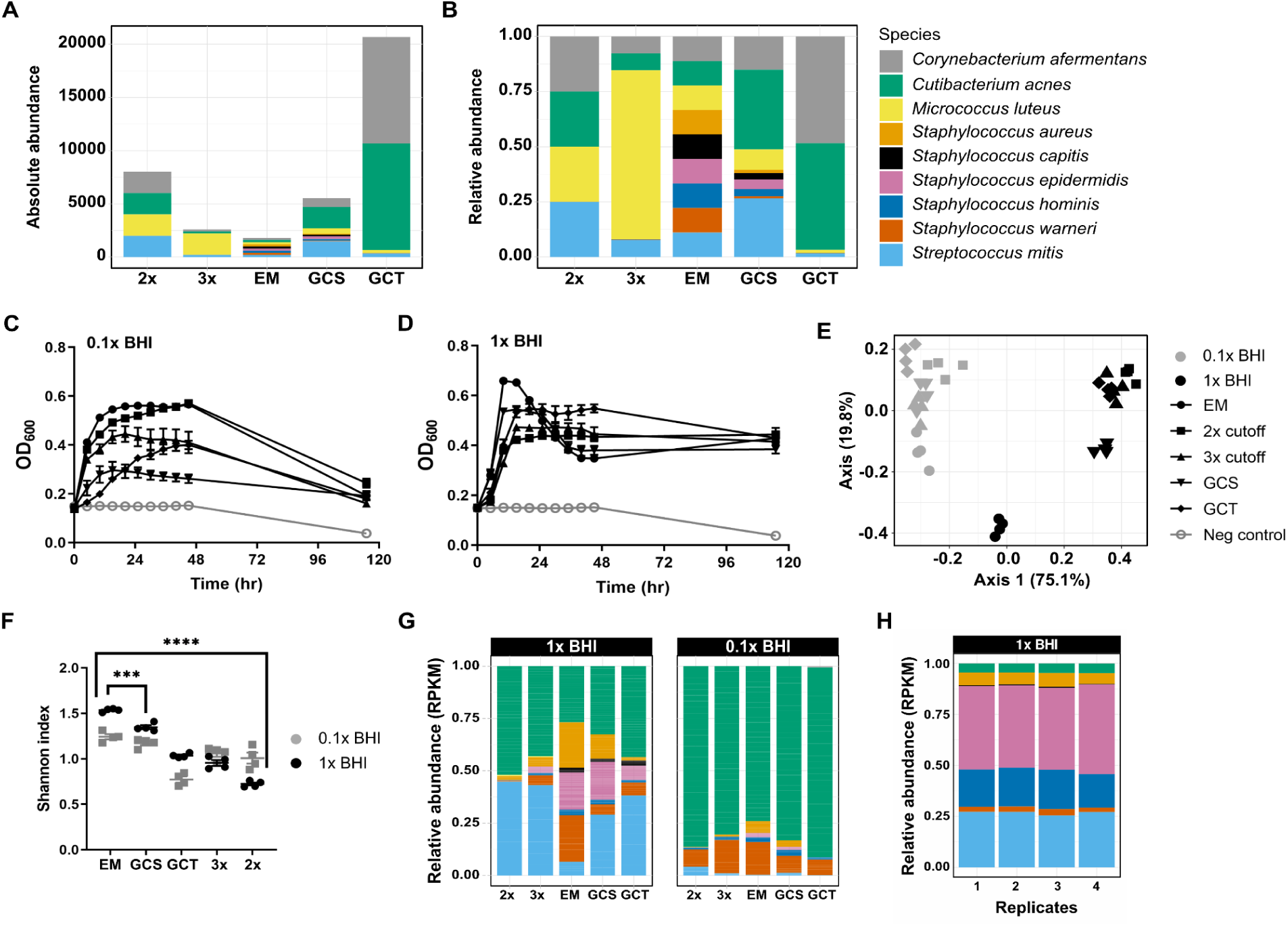
Growth, diversity, and composition of SkinComs with varied starting ratios. A) Absolute abundance, i.e. number of droplets added (0 hr) of each starting ratio. See also Table S1. B) Relative abundance of droplets added (0 hr) of each starting ratio. C) Growth profile of communities from different starting ratios (n = 8 per condition) in 1X BHI and D) in 0.1X BHI. Communities were grown aerobically in 200*μ*l of liquid BHI, non-shaking, at 37 °C. Data are represented as mean ± SEM. E) PCA of Bray-Curtis distances between communities in 1X and 0.1X BHI. F) Shannon diversity index of communities grown in 1X and 0.1X BHI (n = 4 per condition). G) Relative abundance (normalized to RPKM) of communities at each ratio grown in 0.1X and 1X BHI. H) Relative abundance (normalized to RPKM) of GCS communities grown in 1X BHI (n = 4).

We experimented with four distinct starting ratios—2X, 3X, Growth Curve (GC) slope adjusted (GCS), and GC time adjusted (GCT)—and compared them to the standard equally-mixed (EM) configuration. Our goal was to create a more balanced community, potentially mimicking the natural diversity and resilience of the skin microbiome more closely. Upon culturing these adjusted communities under identical conditions to the EM setup, we observed that while the EM community initially showed the highest growth peaks in both diluted (0.1X) and standard concentration (1X) BHI media, all configurations eventually converged to similar final optical densities (OD600) after a 120-hour growth period. (Figure 3C and 3D). Of note, the GCS community had the lowest maximum OD600 in diluted media but the second-highest in standard media, indicating that adjustments based on the growth curve slope might favor resilience or adaptability in nutrient-rich environments. On the other hand, the 2x community, designed with a simple fast/slow grower distinction, had the lowest OD600 in 1X BHI but the second-highest in 0.1X BHI. Diversity analysis revealed significant differences among the communities, with the EM setup maintaining the highest alpha-diversity, followed in descending order by GCS, GCT, 3X, and 2X configurations (Figure 3C). Bray-Curtis dissimilarity analysis, which quantifies the compositional difference between communities, confirmed significant variations between the communities grown in diluted versus standard BHI media (Figure 3D).

In selecting a SkinCom for subsequent experiments, we considered both the alpha-diversity and stable taxonomic composition of the communities. Notably, every community facilitated significant growth of *C. acnes*, the most abundant microbe on human skin. Principal component analysis, using Bray-Curtis distance, revealed that the 1X EM and 1X GCS communities cluster distinctly from the other groups (Figure 3E). In particular, the EM community in 1X BHI had the highest alpha-diversity and contained the highest abundance of *S. aureus* among all communities (Figure 3F). *S. aureus* is a common source of skin infections and has been associated with skin diseases such as atopic dermatitis^14,15,32–35^. Despite its rapid growth to peak OD600 in 1X BHI, the EM community’s OD600 then fell sharply, indicating a swift population decline.

Consequently, we considered the community with the second-highest diversity, i.e. GCS, for downstream experiments. The GCS community showed significantly lower presence of *S. aureus* compared to the EM community, maintained a higher OD600 at the end of the growth period, and showed excellent reproducibility of community composition across replicates (Figure 3H, Supplemental Figure 4). Therefore, our next studies will focus on the GCS community in 1X BHI, given its balanced diversity and stable growth characteristics.

### SkinCom application in murine epicutaneous model

Our ability to fine-tune the composition of the SkinCom opens doors to more complex studies such as those involving *in vivo* models. We therefore investigated if the SkinCom could be successfully incorporated into an existing epicutaneous murine model, aiming to recover both DNA and RNA following skin application *in vivo* to enable researchers to pose functional questions in future studies^37^. The study involved applying the SkinCom onto the dorsal skin of 8 weeks-old female CD1 mice for three days^38,39^. e grouped five mice each to receive the SkinCom with either no *S. aureus*, low *S. aureus* (10^2^ CFU/ml), medium *S. aureus* (10^4^ CFU/ml), or high *S. aureus* (10^6^ CFU/ml). We choose varying concentrations of *S. aureus* cells to demonstrate the suitability of the SkinCom for skin disease research, i.e. atopic dermatitis^14,15,32–35^. Skin microbiome swabs were collected from the dorsal skin and processed for metagenomics and metatranscriptomics library preparations^40^. We successfully recovered DNA and RNA from the SkinCom species and the mice skin microbiome (Figure 4). Additionally, no bacterial growth was detected in collected blood, indicating that the SkinCom patch application did not cause systemic infection (Supplemental Table S2). This result establishes that we can fine-tune the SkinCom inoculum to study effects of defined microbial populations, apply it to an *in vivo* murine model, and recover DNA and RNA from both the SkinCom and resident mouse microflora up to 72 hours post-application.

**Figure 4.**
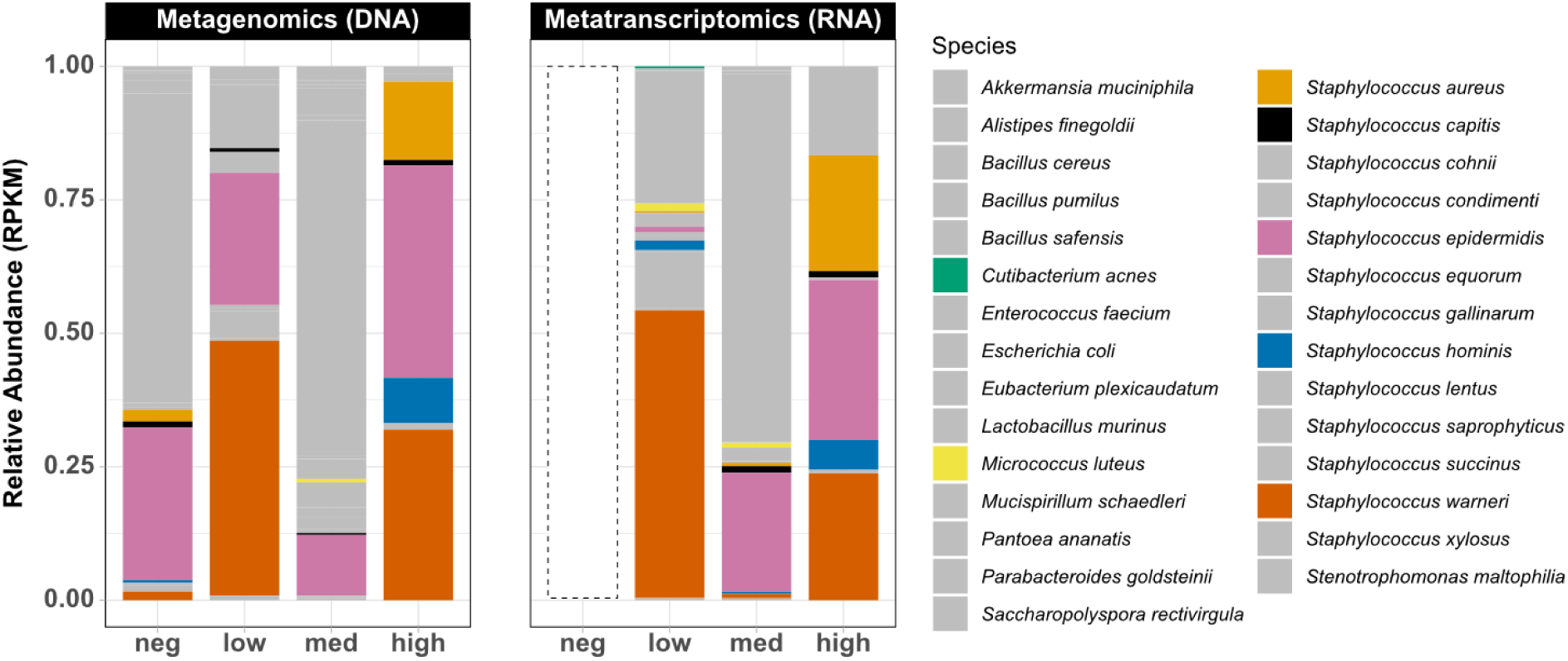
SkinCom application and recovery of DNA and RNA from a murine epicutaneous model. Relative abundances of SkinCom and murine skin microbiome after three days of SkinCom application on CD1 mice (n=5 x 4 groups). Grey represents species not included in the SkinCom; neg = no SkinCom, low = 10^2^ CFU/ml per SkinCom species, med = 10^4^ CFU/ml per SkinCom species, high = 10^6^ CFU/ml per SkinCom species. Dashed box indicates insufficient reads from metatranscriptomics sequencing due to low RNA recovery.

### Effect of cosmetics chemicals on SkinCom

Next, we deployed the GCS community to assess the impact of chemicals commonly found in cosmetics on skin microbes and their resilience to exogenous elements. Investigations into the skin microbiome have highlighted its significance in skin health and dermatological conditions, alongside extensive studies on the effect of skin cosmetics on the microbiome^8,41,42^. However, there is no standardized methodology for evaluating the effects of skin care compounds on the microbiome. We therefore tested the ability of the *in vitro* SkinCom to predict potential impacts of these compounds on *in vivo* skin microbes, potentially providing a rapid and cost-effective approach for the prioritization of new skin care products.

We exposed the GCS SkinCom to four chemicals prevalent in skin and personal hygiene products: sodium lauryl sulfate (SLS, also known as sodium dodecyl sulfate), sodium laureth sulfate (SLES), a rhamnolipid (RL, i.e. RHEANCE®), and creatine (Crt) (Figure 5A)^43–47^. SLS, an anionic detergent known for its cleansing properties in soaps and shampoos, is often used as an irritant in contact dermatitis models^43^. SLES, another anionic detergent, is considered milder than SLS but can still cause skin irritation^44^.

**Figure 5.**
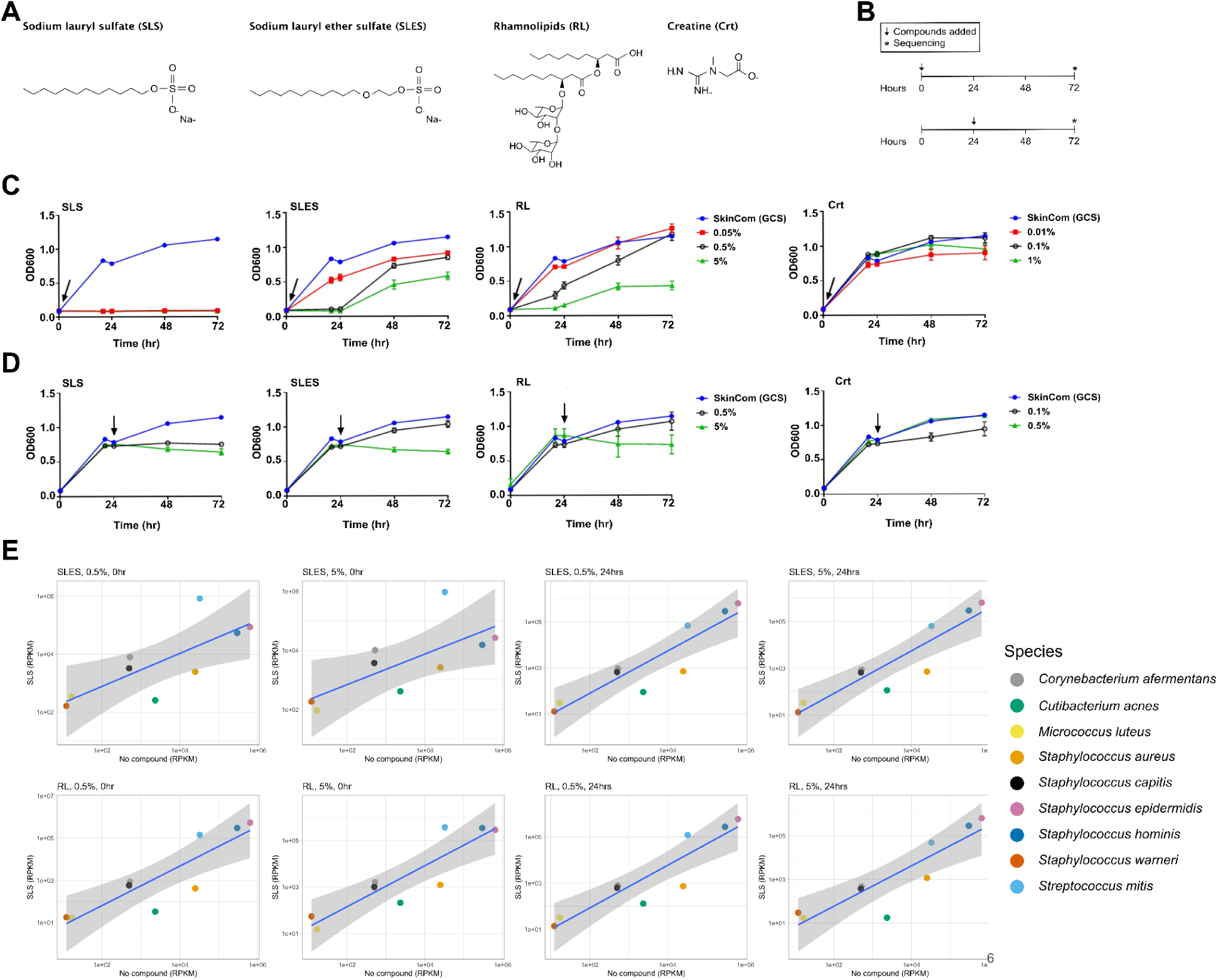
Effect of skin product compounds on SkinCom growth and composition. A) Compounds used in the experiment: Sodium Lauryl Sulfate (SLS), Sodium Lauryl Ether Sulfate (SLES), Rhamnolipids (RL), Creatine (Crt). B) Timeline of the experiment. Compounds were added at either time of inoculation (0 hr) or 24 hrs post-inoculation; the addition of compounds indicated by a red arrow. Compounds were added in different concentrations (w/v) (n = 4 per condition). C) SkinCom growth in the presence of compounds either added immediately upon SkinCom assembly or D) at 24-hr post assembly as measured by OD600. For each plot, the compound addition time is depicted by a black arrow. The growth profile of the SkinCom without additions (SkinCom only; blue line) is provided on each plot for reference. Data are represented as mean ± SEM. E) Changes in SkinCom taxonomic composition in the presence of SLES and RL, as determined by shotgun metagenomic sequencing. RPKM of the indicated compound at 0.5% concentration, added at 0hr or 24hrs, is plotted against RPKM of the SkinCom only (no compound added). The blue line represents a linear regression of the community comparison, with a 95% confidence interval indicated by the shaded gray area. RPKM is averaged between the four replicates for each condition, with error bars representing the standard error of the mean. See also Figure S1.

Rhamnolipids are a class of surfactants produced in various forms by a variety of bacteria, notably *Pseudomonas aeruginosa*^45^. They are typically considered less toxic, more biodegradable, and more environmentally-friendly than detergents such as SLS and SLES^46^. Creatine, a naturally-occurring compound involved in cellular energy homeostasis, is believed to enhance skin cell health and reduce wrinkles when applied topically^47^. To examine these compounds’ effect *in vitro*, we introduced each chemical to the SkinCom in different concentrations (w/v, see Methods), at the start of growth (0 hr) or after 24 hrs, to mimic exposure to an established skin microbiota (Figure 5B). The treated SkinComs were incubated for 72 hrs before undergoing shotgun metagenomics sequencing.

Optical density measurements revealed that all concentrations of SLS inhibited all further microbial growth both at inoculation and 24 hrs post-inoculation, even at the lowest concentration of 0.05%. SLES and RL showed a dose-dependent inhibition of microbial growth when added at inoculation, with SLES demonstrating a stronger inhibitory effect than RL (Figure 5C-D). Both SLES and RL halted further microbial growth with a 5% treatment at 24 hrs, while the 0.5% treatment had only mild effects. Crt treatments did not significantly alter the growth profile of the SkinCom, with slight reductions in 0.01% creatine compared to 0.1% creatine at 24 hrs, but these changes were not statistically significant.

Further investigation focused on the effects of SLES and RL, as these allowed for substantial community growth in a dose-dependent manner at both time points. Through shotgun metagenomics sequencing of the SkinCom at 72 hrs post-treatment, then comparing the reads per kilobase million (RPKM) for communities treated with a compound to the RPKM of untreated communities, we identified microorganisms whose relative abundance changed upon exposure to these compounds. SLES and RL at 0.5% led to a notable shift in community composition, particularly a decrease in *C. acnes* and *S. aureus* (Figure 5E). The alterations were more pronounced for SLES added at 24hrs compared to at 0hr. On the other hand, the decrease in *C. acnes* and *S. aureus* were consistent regardless of the timing for RL application (0 hr or 24 hrs). The composition changes observed with RL were similar to the changes observed with SLS and SLES (Supplemental Figure 1).

### Responses of SkinCom and human skin microbiome to cosmetic compounds

To assess how the outcomes from the *in vitro* trial align to observations of the human skin microbiome, we conducted a human-subject study to investigate the impact of cosmetic ingredients, specifically RL and SLES, on the skin microbiome of the forehead. This trial aimed to gather preliminary data and offer initial insights into SkinCom’s relevance to human skin. Our cohort consisted of 13 individuals, of which five applied SLES and eight applied RL to one side of their forehead for 30 seconds, twice a day (see Materials & Methods). As a control, subjects additionally applied a cotton round soaked with water to the opposite side of their forehead. Swabs were collected from the forehead on days 4 and 7 for shotgun metagenomics analysis, processed following established protocols^34,40^.

From the *in vitro* study, we conducted a differential abundance analysis to determine the changes in abundance of each SkinCom species in relation to a reference species^48^. We observed a significant reduction in *C. acnes, S. aureus,* and *S. mitis* (t-statistic < 0 and p-value < 0.05), and a significant increase in *S. hominis* and *S. warneri* from SLES treatment (t-statistic > 0 and p-value < 0.05 (Supplemental Table S3). For samples that received RL treatments, we observed a significant reduction in *C. acnes*, *S. aureus*, *S. capitis*, and significant increase in *S. epidermidis*, *S. hominis*, and *S. warneri* in RL treatment (Supplemental Table S4).

For the human study, principal component analysis revealed individual variances, consistent with existing literature^10,17,18^(Supplemental Figure 3). For the SLES cohort, differential abundance showed a reduction in *C. acnes, M. luteus, S. aureus, S. epidermidis,* and *S. warneri* on day 4 (Supplemental Table S5), and a reduction in *C. acnes* and *S. mitis* on day 7 (Supplemental Table S6). Meanwhile in the RL cohort, differential abundance showed a reduction in *C. acnes, S. aureus,* and *S. warneri* on day 4 (Supplemental Table S7), and a reduction in *C. acnes*, *M. luteus*, *S. hominis*, and *S. mitis* on day 7 (Supplemental Table S8). Note that the only statistically significant change observed was for *S. epidermidis* in the day 7 SLES cohort. No other changes were statistically significant.

Remarkably, an average of 65.6% of the shifts observed *in vitro* were also seen in the human subject trial (Figure 6), suggesting that the SkinCom can capture the majority of responses to perturbation by topical exposure in the skin microbiome. The breakdown is as follows: 62.5% agreement in the SLES group on day 4, 87.5% agreement in the SLES group on day 7, 75% agreement in the RL group on day 4, and 37.5% agreement in the RL group on day 7.

**Figure 6.**
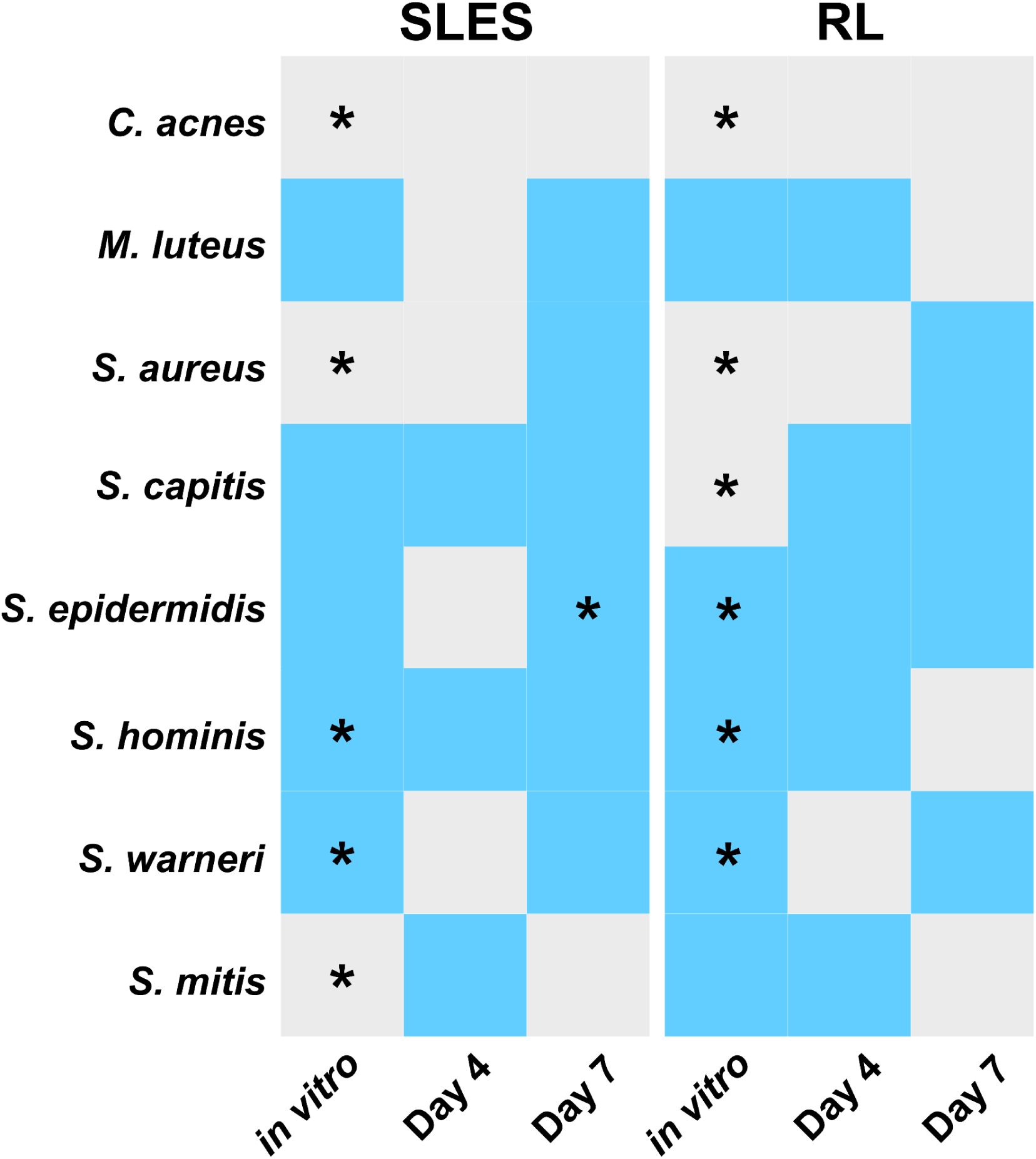
Summary of average differential abundance analyses between SkinCom and human trial. Changes after compound applications *in vitro* and *in vivo* (human trial: Day 4 or Day 7). For the *in vitro* experiment, SLES and RL were tested at 0.5% and 5%, for 3 days. For the human trial, both SLES and RL were tested at 3% for 4 or 7 days. Skin swabs were collected prior to chemical applications and 4 and 7 days post chemical applications. Grey: average differential abundances of samples with control is larger than that of chemical treatment; pre-treatment species are more abundant than post-treatment (t statistics is negative). Blue: average differential abundances of samples with chemical treatment is larger than that of control; post-treatment species are more abundant than pre-treatment (t statistics is positive) * indicates p < 0.05.

## Discussion

The skin microbiome is a vital determinant for skin health, but its study has been proven complicated by low abundance of microbial cells, its dynamic nature, and individual variability. In response, we developed a model synthetic bacterial community (SkinCom) to facilitate reproducible research into the skin microbiome. We optimized the SkinCom for composition, diversity, and reproducibility *in vitro* before applying it to an epicutaneous infection murine model. This application demonstrated the model’s utility in recovering metagenomic and metatranscriptomic data three days post-application, underscoring its value in addressing *in vivo* skin microbiome and pathogen studies. Furthermore, we explored the effects of common cosmetic compounds on the SkinCom and compared these *in vitro* results with those from a human subject trial. This comparison showed that 65.6% of the microbial changes observed *in vitro* were reflected in the human study, suggesting that the SkinCom effectively models skin microbiome responses to topical perturbations.

The SkinCom consists of nine bacterial species spanning five genera, representing the most prevalent species found on the human skin. Using the approaches we developed, the nine-member consortium could readily be expanded to represent body site-specific skin microbiota. Furthermore, our methods allow for precise identification of each community member through shotgun metagenomic sequencing.

Interestingly, *C. acnes*, typically anaerobic, proliferate in oxic conditions when co-cultured with aerobic bacteria, likely due to oxygen scavenging by the latter. We hypothesize that the incubating conditions allowed the microbes to respire molecular oxygen and thus create an oxygen gradient within each well, resulting in reduced oxygen concentrations at the bottom of the well. This oxygen-scavenging by other microorganisms permits the growth of the anaerobic *C. acnes*^49^. This phenomenon and the high alpha-diversity achieved in a nutrient-rich medium hint at the natural adaptability of skin microbes to rich environments found within skin microenvironments like hair follicles and sebaceous glands^50,51^.

Our ability to adjust community proportions in the SkinCom enables the simulation of varied skin microbiome profiles. This feature is particularly useful for studying pathogens such as *S. aureus,* associated with skin conditions such as atopic dermatitis^14,15,32–35^, and assessing interventions designed at modulating its abundance. Reducing the amount of *S. aureus* starting inoculum based on its growth rate (GCS) resulted in a reduced relative abundance of this pathogen in the final community, akin to a healthy skin microbiome (Figure 3G). Moreover, our ability to recover both DNA and RNA from the synthetic community is uncommon due to colonization resistance^52,53^, but has been reported^54^. This allows for future investigations of microbe-microbe interactions in combination with metagenomics or metatranscriptomics tools.

SLS, SLES, and RL are molecules routinely included in skin care products for their detergent or surfactant properties. Treatment of the SkinCom with the detergent SLS prevented all microbial growth even in very small concentrations in agreement with previous work on the harsh effects of SLS^55,56^. Treatment with SLES also inhibited community growth in our *in vitro* setting, although to a lesser extent than SLS. This indicates that although SLES is considered a gentler compound than SLS, it still has an inhibitory effect on microbial growth. All three compounds, i.e. SLS, SLES, and RL, caused similar changes in community composition profiles (Figure 5E, Supplemental Figure 1), most notably a decrease in the relative abundance of *S. aureus* and *C. acnes* and an increase in *S. warneri.* Strikingly, *C. acnes* can be both a commensal and pathogenic member of the skin community, depending on the virulence factors carried by the individual strains^57,58^ and further investigations are needed to confirm the potential effect of Crt on skin health and the microbiome.

The utility for the SkinCom in studying human microbiome responses was established by comparing *in vitro* SkinCom responses to *in vivo* skin microbiome changes after compound applications. While we observed over 65.6% agreement between these two studies overall, i.e. *in vitro* and *in vivo,* it is important to note that these preliminary studies are of low statistical power (n=13). Moreover, due to such different experimental designs, results cannot be used interchangeably. Some explanations for discrepancies in absolute abundance changes between the two study designs are: the duration of the experiments, compound dosage, species and strain resolution, and most importantly the host and environmental exposure. The *in vitro* study measured species composition after three days of growth in the presence of compounds, while the human subject trial measured results at day 4 or day 7. Another explanation for some discrepancies between the *in vitro* and the *in vivo* study is likely due to the varying degree of environmental exposure that human subjects face during the 7-day trial period, as reports have shown that in addition to cosmetics compounds, day-to-day environmental exposures significantly contribute to changes in the skin microbiome composition^6–8^. Additionally, even though the skin microbiome is generally considered stable over time^10,18^ natural temporal variations can occur over the course of the study. These variations can also include strain-level differences, which are currently not captured by standard shotgun metagenomic approaches^59^. However, the main reason for the observed differences between the *in vitro* and the *in vivo* study is very likely due to the host actively influencing the microbiome and the fact that the *in vitro* study was carried out in rich medium, providing close to optimal conditions for all bacteria. While these studies are thus not directly comparable given these significant differences, the SkinCom provides a highly reproducible tool for the study of the skin microbiota and can provide initial hypotheses to be tested in human trials. Furthermore, we recommend that differential abundance analysis is conducted for future analyses involving microbial abundance changes due to its ground truth value built into the dataset, either by the organism changing the least or controls such as synthetic DNA^60,61^.

In summary, the SkinCom presented here can serve as a simplified model of the cutaneous microbiome, consisting of both aerobic and anaerobic microorganisms typically found on human skin and applicable in both *in vitro* and *in vivo* settings. The high reproducibility of the SkinCom renders it a suitable tool for studying the response of the skin microbiota to perturbations, e.g. the effect of skincare compounds. Moreover, the SkinCom can be applied in a murine model to study disease progression along with corresponding changes in microbial load. Our results further present the first synthetic community for human skin consisting of both aerobic and anaerobic skin bacteria—together spanning five genera. We anticipate the SkinCom and modifications thereof to be of broad use for the study of the human skin microbiome, such as those investigating genomic, functional, or metabolite dynamics.

## Acknowledgements

The authors would like to thank Peter Lersch and the EVONIK team for the opportunity to investigate the effects of chemical compounds on the skin microbiome. Furthermore, we would like to thank our collaborators Anna DiNardo and Kana Kuroki (UCSD) for performing a pilot study involving a murine model. We are also thankful for all study volunteers of the human trial and for Stephanie Flores Ramos (UCSD) for early work related to SkinCom assembly. Additionally, the authors would like to thank the past and present members of the Zengler Lab for their experimental contribution, lively discussion, and critical evaluation of this manuscript. This work was partly supported by Evonik Industries AG and the UC San Diego Center for Microbiome Innovation.

## Author contributions

KZ, JC, and CM conceptualized the study. AL, DT, JC, FA, AK, CM, and AH contributed to acquisition of data. AL, DT, JC, FA, and AK contributed to data analysis. AL, DT, JC, FA, AK, CM, VN, and KZ contributed to data interpretation. AL, DT, JC, FA, AK, VN, and KZ drafted the manuscript. AL, DT, JC, FA, AK, VN, and KZ revised the manuscript.

### Funding

KZ and VN acquired funding for the study. All authors read and approved the final version of the manuscript and had access to all the data in the study.

## Declaration of interests

The authors declare no competing interests.

*Corynebacterium afermentans* is used as the denominator for the log ratio test. See also Table S3-8.

## STAR METHODS

Detailed methods are provided in the online version of this paper and include the following:

- KEY RESOURCES TABLE
- RESOURCE AVAILABILITY

○ Lead contact
○ Materials availability
○ Data and code availability
- EXPERIMENTAL MODEL AND SUBJECT DETAILS

○ Individual strain growth conditions
○ Community assembly with CellenONE liquid printer
○ Community growth conditions
○ Subject recruitment and sample collection
- METHOD DETAILS

○ Community growth with cosmetic compounds
○ Shotgun metagenomics library preparation and sequencing
○ Metagenomics sequencing analysis
○ Differential abundance analysis
○ SkinCom application in murine epicutaneous model
- QUANTIFICATION AND STATISTICAL ANALYSIS

## STAR METHODS

### KEY RESOURCES TABLE

| REAGENT or RESOURCE | SOURCE | IDENTIFIER |
| --- | --- | --- |
| Bacterial strains | Source | Medium |
| *Corynebacterium*<br>*afermentans* (ATCC<br>51403, strain CIP 103499<br>[LCDC 88199]) | ATCC | Brain heart infusion medium |
| *Cutibacterium acnes*<br>(ATCC KPA171202) | ATCC | Brain heart infusion medium |
| *Micrococcus luteus* (ATCC<br>4698) | ATCC | Brain heart infusion medium |
| *Staphylococcus aureus*<br>(ATCC 35556, strain<br>SA113) | ATCC | Brain heart infusion medium |
| *Staphylococcus capitis*<br>(ATCC 27840, strain LK<br>499) | ATCC | Brain heart infusion medium |
| *Staphylococcus epidermidis* (ATCC 12228) | ATCC | Brain heart infusion medium |
| *Staphylococcus hominis* (ATCC 27844, strain DM 122) | ATCC | Brain heart infusion medium |
| *Staphylococcus warneri* (ATCC 27836, strain AW 25) | ATCC | Brain heart infusion medium |
| *Streptococcus mitis* (ATCC 49456, strain NCTC 12261). | ATCC | Brain heart infusion medium |

#### Resource availability

All requests for information regarding reagents and resources should be directed to the lead contact and will be fulfilled by the lead contact.

#### Lead contact

Further information and requests for resources and reagents should be directed to and will be fulfilled by Dr. Karsten Zengler.

#### Materials availability

This study did not generate new unique reagents.

#### Data and code availability

##### In vitro experiments

For metagenomics analysis, reads from raw FASTQ files were processed with Trimmomatic (v0.36) to remove adapters and trim low-quality base calls using the parameters “ILLUMINACLIP:NexteraPE-PE.fa:2:30:10 LEADING:10 TRAILING:10 SLIDINGWINDOW:4:15 MINLEN:36”^62^. Trimmed reads were first aligned to the minikraken2 database to exclude possible contamination from non-community organisms^63^. Following this, trimmed reads were aligned to a custom database of community strain genomes using bowtie2 (v2.2.3) using default settings and read counts were transformed into RPKM^64^. Alpha-and beta-diversity was calculated in phyloseq^65^. Beta-diversity plots were generated using phyloseq. Figures 2A-C, 3C-F, and 5C-D were generated using GraphPad Prism 7 software^66^. The code used to process and analyze sequencing data is available on Github (https://github.com/jkccoker/Skin_synthetic_community/tree/main). New sequences have been deposited to the Sequence Read Archive with submission numbers SUB14340232 and SUB14340534 and BioProject number PRJNA1091705.

##### In vivo experiments

Reads from raw FASTQ files were processed with trim_galore (version 0.6.4) to remove adapters^67^. Next, trimmed reads were aligned to a custom reference genome extracted from the Web of Life database^68^ using bowtie2 (v2.2.3) using default settings and read counts were transformed into RPKM^64^. All other plots were generated using R Studio Version 2023.09.1+494. R packages necessary for analysis and visualization include tidyverse, dplyr, ggplot2, readr, stringr, colorspace, RColorBrewer, magrittr, ggpubr, vegan, plyr. ^69–79^ The code used to process and analyze sequencing data is available on Github (https://github.com/alekbua/SkinCom). New sequences have been deposited to the Sequence Read Archive with submission numbers SUB14342487 and SUB14347828 and BioProject number PRJNA1091705. Any additional information required to reanalyze the data reported in this paper is available from the lead contact upon request.

### EXPERIMENTAL DETAILS AND SUBJECT DETAILS

#### Bacteria species selection and growth conditions

From Byrd 2018, Table 1 shows the top ten species of bacteria found in the four microenvironments of the human skin <u>(Byrd et al. 2018)</u>. We selected overlapping species from these four sites, and reduced the number to as few as possible, to create our nine-member SkinCom. All strains were purchased from ATCC: *Corynebacterium afermentans* (ATCC 51403, strain CIP 103499 [LCDC 88199]), *Cutibacterium acnes* (ATCC KPA171202), *Micrococcus luteus* (ATCC 4698), *Staphylococcus aureus* (ATCC 35556, strain SA113)*, Staphylococcus capitis (ATCC 27840, strain LK 499), Staphylococcus epidermidis (*ATCC 12228)*, Staphylococcus hominis* (ATCC 27844, strain DM 122), *Staphylococcus warneri* (ATCC 27836, strain AW 25), and *Streptococcus mitis* (ATCC 49456, strain NCTC 12261). Strains were streaked out on Brain-Heart Infusion (BHI; Millipore Sigma 53286) agar plates prior to making glycerol stocks to confirm purity of the individual microorganisms. Individual strains were cultured in sterile BHI broth at 37 °C, without shaking. 1X BHI was made as directed by manufacturer instructions (37 g BHI powder in 1 L water). 0.1X and 0.2X BHI were obtained by diluting 1X BHI with sterile water. For anaerobic growth, BHI broth was anoxified by bubbling with N_2_ and CO_2_ (80%, 20%). The container was then sealed and the headspace exchanged with N2 and CO2. L-cysteine (Sigma Aldrich 168149) was added to a final concentration of 2 mM directly to the containers before culturing. For *C. acnes* cultures, hemin and vitamin K (VWR 75803-006) were added to final concentrations of 2.5 ug/mL and 0.5 ug/mL, respectively, directly before culturing.

#### Community assembly with CellenONE liquid printer

Each bacteria species was grown to mid-log phase and diluted in BHI to an OD600 of 0.07 after subtracting the blank reading. The diluted strains were loaded into a 384-well “probe” plate, one strain per well. The CellenONE X1 liquid printer (SCIENION US Inc., Phoenix, AZ) was programmed to pick up 30*μ*l from a well of the probe plate using a Piezo Dispense Capillary (PDC) and dispense the specific number of drops (see Table 1) in the appropriate wells of a 96-well “target” plate, which was preloaded with 200*μ*l BHI/well. Prior to dispensing into destination wells, each aliquot was confirmed by eye to have 4-5 cells/drop using the CellenONE camera. OD600 readings were then monitored with the spectrophotometer (Molecular Devices SpectraMax M3 Multi-Mode Microplate Reader, VWR, cat # 89429-536). The PDC was cleaned between isolates by flushing the PDC interior with 0.5mL water. 200 drops of BHI were added to negative control wells as the last step in each experimental setup, to ensure no contamination occurred due to incomplete flushing of the PDC between strains.

#### Community growth conditions

Communities were grown in 200*μ*l of BHI broth in 96-well plates, at 37 °C, without shaking. To minimize evaporative loss, plates were set on 4 100 mm-diameter Petri dishes (2 stacks of 2 dishes) filled with ∼20 mL water each to generate a humid environment around the plates. To prevent condensation of the plate lid which would interfere with spectrophotometric readings, each plate lid was coated with 3 mL of an aqueous solution with 20% ethanol and 0.01% Triton X-100 (Sigma, cat # X100-100ML). Excess liquid was removed after 30 seconds and the lid was allowed to air-dry for 30 minutes under a UV light for sterilization. Under anaerobic conditions, plates were incubated in a vinyl anaerobic chamber (Coy Lab Products) with an atmosphere of N2/CO2/H2 (85%/10%/5%). Plates were incubated at 37 °C without shaking and OD600 readings were taken in a Molecular Devices SpectraMax i3 spectrophotometer with a StakMax Microplate Handling System.

#### Subject recruitment and sample collection

Thirteen healthy adults (9 male, 4 females) were recruited to donate samples. All individuals signed a written informed consent in accordance with the sampling procedure approved by the University of California, San Diego Institutional Review Board (Approval 801694). Subjects with known allergies to cosmetics were excluded as stated in the IRB. Subjects were instructed to avoid applying topical products onto the face during the trial period. Two collections were performed from each skin spot for three time points: Days 0, 4, and 7. Sampling was performed by applying pressure to a 2 x 2 cm area of one side of the forehead in a circular motion for 60 seconds with sterile cotton swabs (Fisher #22-029-630) pre-moistened with a swabbing solution (Tris-EDTA (TE) buffer + 0.5% Tween-20 + 1% TritonX-100). Sample locations were noted and swabs were placed in cryotubes with in 700*μ*l DNA/RNA Shield Buffer (Zymo Research # R1100-50) at -80 °C until sample extraction time. For skincare compound applications, subjects applied a cotton round saturated with either SLES solution (3% in water) or RL solution (3% in water) to the treatment side of their forehead for 30 seconds, twice a day. We processed skin swabs by extracting DNA and RNA using the ZymoBIOMICS DNA/RNA miniprep kit (cat# R2002), and prepared metagenomics library as previously described.

### METHOD DETAILS

#### Community growth with cosmetic compounds

Sodium Lauryl Sulfate (SLS), Sodium Lauryl Ether Sulfate (SLES), Rhamnolipid (RL), and creatine compounds were obtained from Evonik Industries (https://corporate.evonik.com/en). For compounds added at time of inoculation, compounds were diluted to the indicated concentration (w/v) in 1X BHI and loaded into the target plate prior to community spotting, 150*μ*l/well. Communities receiving compounds at 24 hrs post-inoculation were spotted into plain 1X BHI and compounds were diluted to 4X the indicated concentration in 1X BHI and 50*μ*l were added to the appropriate wells after 24 hrs of incubation. 50 *μ*l of plain BHI were added to communities that received compounds at time of inoculation. All compound solutions were sterilized by syringe-filtering across a 0.22 um filter before addition. Community plates were grown oxically for days. Plates were then stored at -20 °C until processing for sequencing.

#### Shotgun metagenomics library preparation and sequencing

DNA was extracted from community samples using a Qiagen DNeasy PowerSoil Pro Kit (cat # 47016) according to manufacturer’s instructions with the following noted change. Samples were heated for 10 min at 100 °C after addition of lysis buffer and prior to vortexing. Following extraction, DNA was quantified with Qubit dsDNA, high sensitivity (ThermoFisher cat # Q32851) and normalized to 0.2 ng/uL. Shotgun metagenomic sequencing libraries were prepared using the Nextera XT DNA Library kit with 1 ng DNA input, according to manufacturer’s instructions (Illumina cat # FC-131-1096 and FC-131-2001). Libraries were quantified using a Qubit as indicated above and normalized to 2 ng/*μ*l for sequencing. Libraries were sequenced on an Illumina MiSeq platform with a paired-end 150 V2 kit (Figure 2, 3, and 5) or a paired-end NovaSeq PE 100 V2 kit (Figure 4 and 6).

#### Metagenomics sequencing analysis

Metagenomics reads on day 0 and day 4 were trimmed using Trim Galore v0.6.10^66^ paired-end mode with default parameters and aligned to the reference database Web of Life (WoL)^80^ using bowtie2 v2.2.5^63^. Taxon count tables were generated from alignment using Woltka v0.1.5^67^.

#### Differential abundance analysis

We conducted differential abundance analyses on the non-rarefied metagenomics data obtained from the *in vitro* SkinCom with cosmetic compounds and human subject trials using Songbird v. 1.0.4 through QIIME2 v. 2020.6.0^48^, with parameters *p-min-sample-count* and *p-min-feature-count* set to 0 due to low microbial counts in skin samples. Rank plots were generated using Qurro v0.8.0^60^. Code used to process differentials is available at Github (https://github.com/sherlyn99/skincom_analysis). Paired t-tests were conducted to assess the change in abundance of each taxon at two time points: before the compound application and on day four, as well as before the application and on day seven.

#### SkinCom application in murine epicutaneous model

Animal experiments was conducted in accordance with the rules and regulations of the Institutional Animal Care and Use Committee, which was approved by the UC San Diego IRB protocol S00227M. Mice were housed in filter-top cages with regulated environmental conditions (20-22 °C, 30-70% relative humidity, 12 h light/12h dark cycle). The SkinCom was constructed in 1X BHI following the GCS ratio previously described^31^ with no (control/mock infected, containing sterile BHI) or the final bacterial load of 10^2^ CFU/ml (low bacterial load), 10^4^ CFU/ml (medium bacterial load), and 10^6^ CFU/ml (high bacterial load). After three days of incubation, cultures were pelleted and spotted onto 2 x 2 cm patch of sterile agar, and affixed onto the chemically depilated area (Nair, USA) of eight-week-old female CD1 mice (Charles River Laboratories, n=5 mice/group) as described previously (Nizet et al., 2001; Hirose et al 2021). Briefly, the mice were shaved under isoflurane sedation 24 h prior to SkinCom implementation. Under sedation, the SkinCom agar patch was securely fastened with sterile gauze and a transparent film dressing (3M Tegaderm) on the shaved skin. To assess the systemic spread of infection 72 h post SkinCom patch application, the heparinized blood was collected submandibularly and serially diluted on BHI agar plates. Sequentially, animals were euthanized by CO_2_ inhalation and skin swabs were collected in accordance with the sample collection guidelines previously outlined for DNA/RNA extractions associated with metagenomics and transcriptomics analysis^33,40^.

#### Quantification and Statistical Analysis

Several statistical methods were conducted to assess differences, correlations, and associations between groups. In Figure 1, the mean of 4 replicates are shown, along with the error bars showing the standard error of the mean. The same statistical analyses were run for figures 2 and 3. For growth curves, 8 replicates per condition were averaged and the standard error of the mean is demonstrated by the error bars. For diversity indices, 4 replicates per media condition were averaged and calculated for alpha- and beta-diversity using phyloseq^65^. The statistical significance of the Shannon diversity index was calculated with the Student’s t-test, p = <0.0001, while a PERMOANOVA test, p = 0.032, was calculated to determine the statistical variation of Bray-Curtis distance. For relative abundance plots, raw read counts from metagenomics sequencing (n=4) were normalized by reads per kilomillion base pairs (RPKM). PCA from each condition was plotted using the Bray-Curtis distances. For Figure 4, aligned reads were calculated for genome coverage (cite) and coverage below 0.02 was filtered out. All reads were normalized by RPKM as previously described^64^. Each bar represents 5 mice receiving the same treatment, residing in the same cage. For figure 5, the standard error of the mean (error bars) was calculated for each of the conditions (4 replicates each). Linear regression with a 95% confidence interval was utilized to determine changes in SkinCom composition in the presence of SLES and RL (n=4). For figure 6, the t-statistic was calculated to analyze the differential abundances of various samples. The statistical significance of the analysis was determined with p < 0.05.

## Supplemental Figures

**Supplemental Figure 1.**
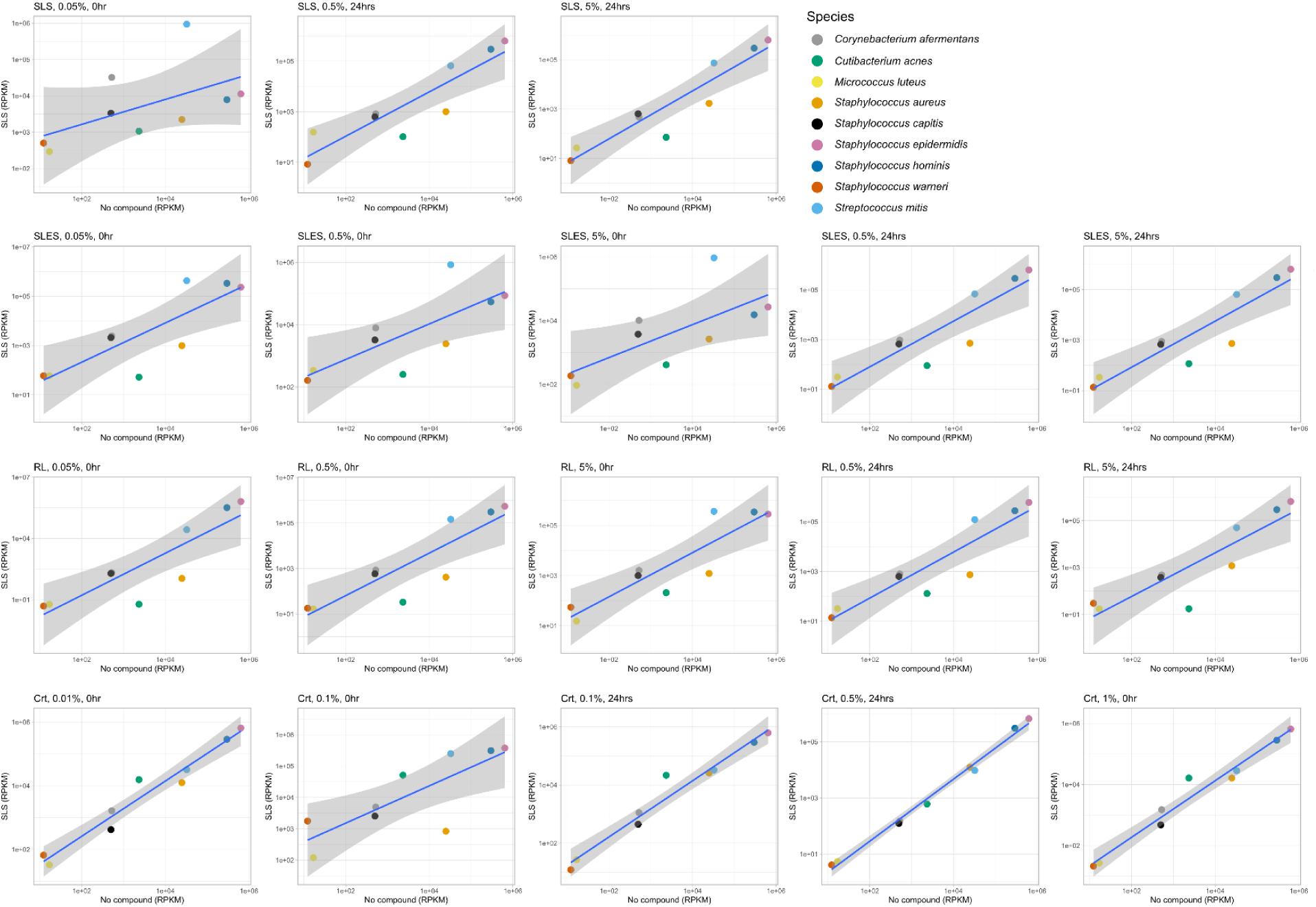
Changes in community composition in the presence of skin product compounds. RPKM of the compound, concentration, and time of compound addition (as indicated by each plot title) is plotted against RPKM of the community alone. The gray dotted line represents no change between communities (x = y). The blue line represents a linear regression of the community comparison, with a 95% confidence interval indicated by the shaded gray area. RPKM is averaged between the 4 replicates for each condition, with error bars representing the standard error of the mean.

**Supplemental Figure 2.**
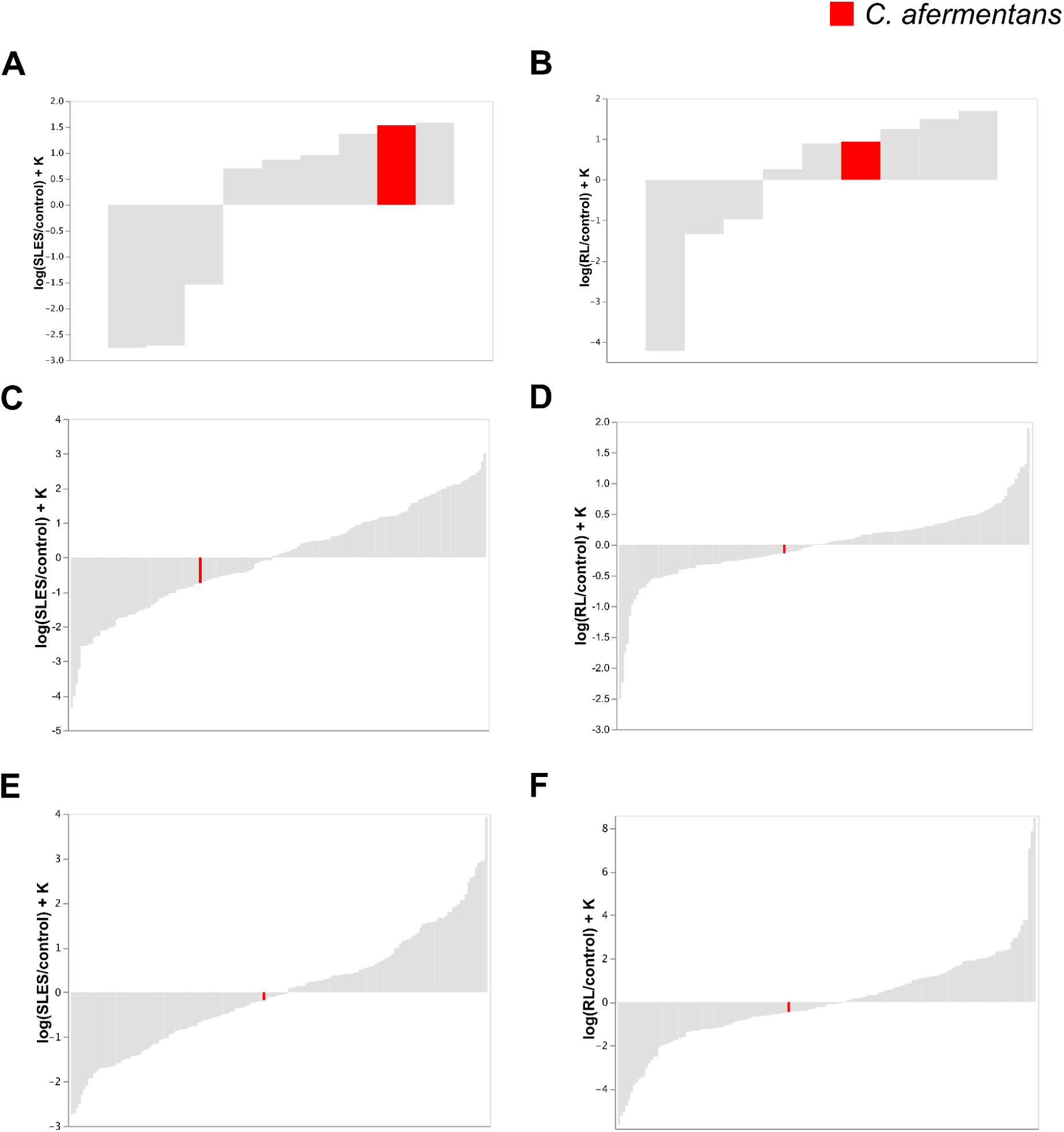
Rank plots of 9 SkinCom strains from differential abundance analysis. Relative differentials estimated using Songbird (QIIME2, v2020.6). The rank of the chosen reference species, *C. afermentans,* was highlighted in rank plots generated from (A) *in-vitro* SkinCom SLES and control samples; (B) *in-vitro* SkinCom RL and control samples; (C) clinical trials day 4 SLES and control samples; (D) clinical trials day 4 RL and control samples; (E) clinical trials day 7 SLES and control samples; (F) clinical trials day 7 RL and control samples. Grey represents all species present in the sample. Red represents *C. afermentans*.

**Supplemental Figure 3.**
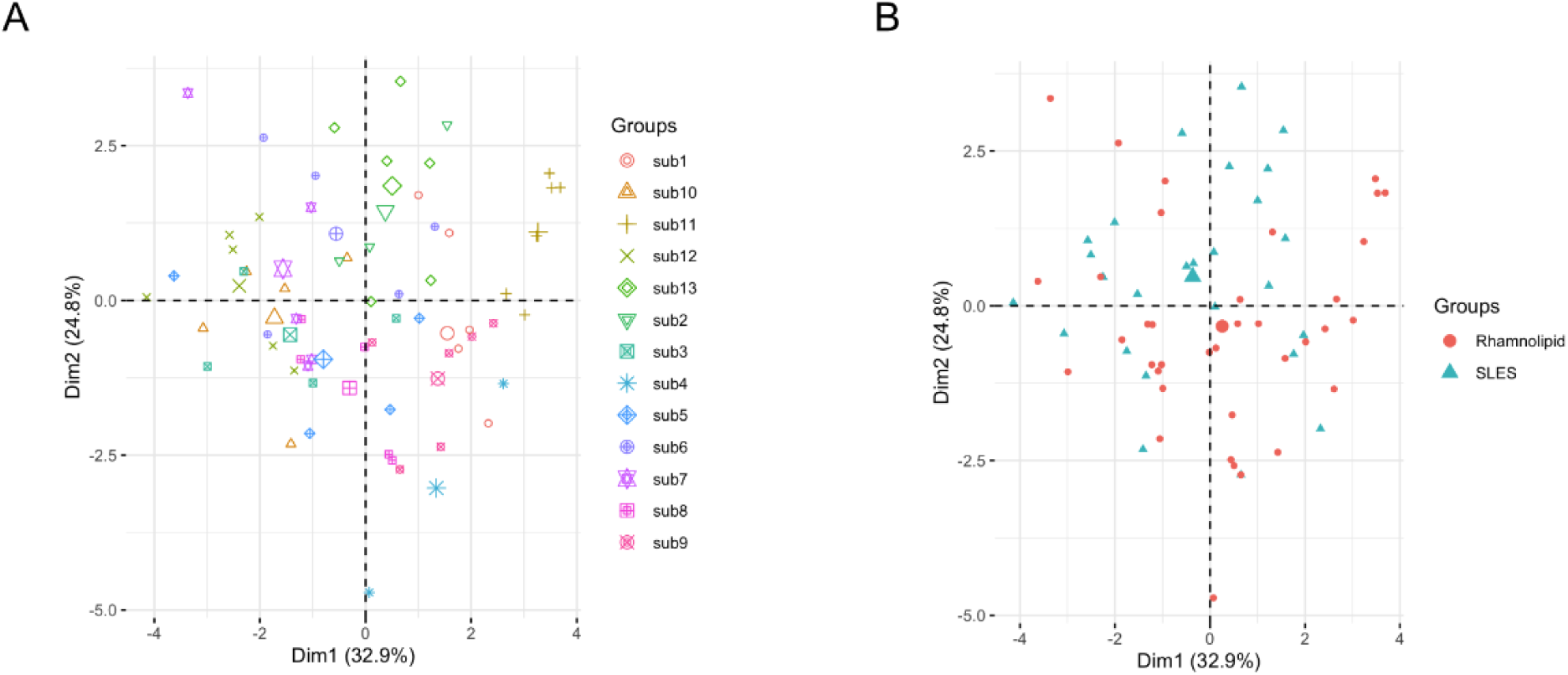
Principal Component Analysis (PCA) based on subjects (A) and chemical treatments (B). Metagenomics sequencing results were processed by trimming (trim_galore version 0.6.4_dev), concatenating, and alignment to the Web of Life database (bowtie2 version 2.3.5.1). PCA analysis by subjects (sub1-13) show that samples from the same individual cluster together, confirming previous skin microbiome findings. Large symbols represent the average between samples from the same group.

**Supplemental Figure 4.**
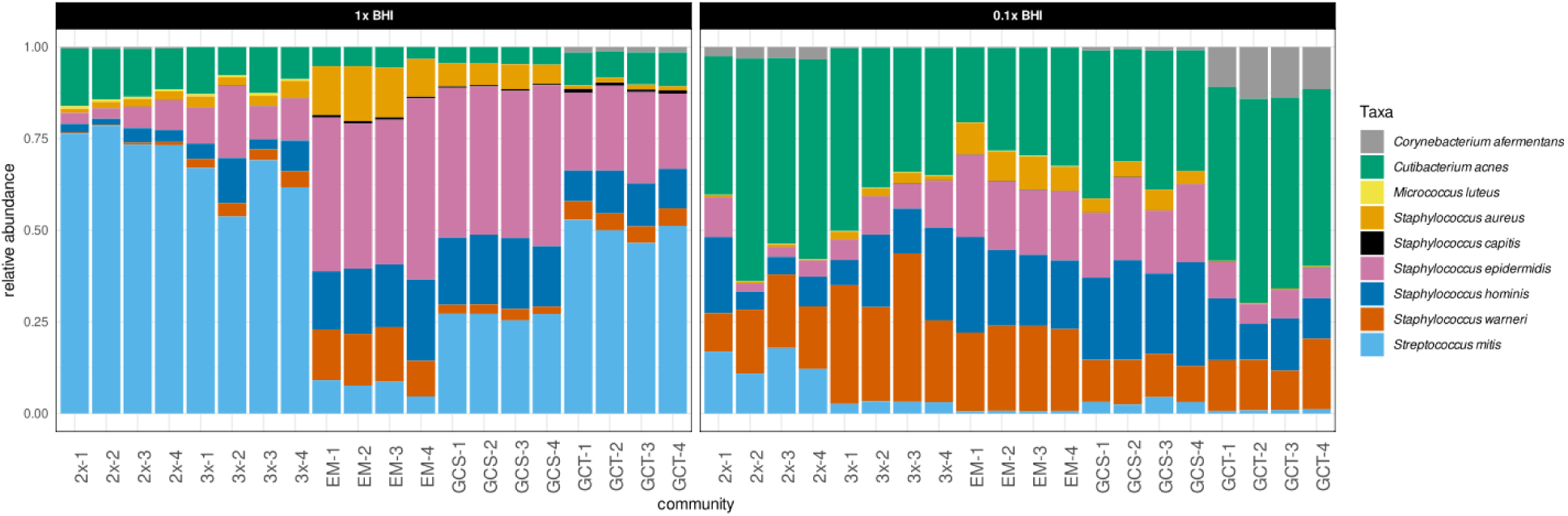
All replicates of all the community combinations and media compositions tested for achieving the optimal SkinCom. Legend is consistent with Figure 3. Two media were tested (1X BHI and 0.1X BHI). n=4 for each condition.

## Supplemental Tables

**Table S1.** Starting amounts of skin microorganisms in five different community ratios, Related to Figure 3. GC = growth curve.

| Organism | Equal mix | 2x cutoff | 3x cutoff | GC slope adjusted | GC time adjusted |
| --- | --- | --- | --- | --- | --- |
| *C. afermentans* | 200 | 2000 | 200 | 835 | 10000 |
| *C. acnes* | 200 | 2000 | 200 | 2000 | 10000 |
| *M. luteus* | 200 | 2000 | 2000 | 510 | 304 |
| *S. aureus* | 200 | 2 | 2 | 83 | 3 |
| *S. capitis* | 200 | 2 | 2 | 161 | 27 |
| *S. epidermidis* | 200 | 2 | 2 | 242 | 7 |
| *S. hominis* | 200 | 2 | 2 | 178 | 9 |
| *S. warneri* | 200 | 2 | 2 | 54 | 6 |
| *S. mitis* | 200 | 2000 | 200 | 1472 | 319 |

**Table S2.** Bacterial detection in heparinized blood. Heparinized blood was collected submandibularly from the epicutaneous murine (n=5 per group), serially diluted in PBS, and spotted on BHI agar plates, then incubated at 37°C for 3 days.

| Group | No SkinCom | Low SkinCom (10<sup>2</sup> CFU/ml) | Medium SkinCom (10<sup>4</sup> CFU/ml) | High SkinCom (10<sup>6</sup> CFU/ml) |
| --- | --- | --- | --- | --- |
| Bacterial colony detection | no | no | no | no |

**Table S3.** Statistics for 24h, *in-vitro* SkinCom, SLES vs Control, Related to Figure 6.

|  | **t statistics** | **p value** | **upperCI** | **lowerCI** |
| --- | --- | --- | --- | --- |
| **log(*C.acnes/C.afermentans*)** | -4.546796 | 0.002646 | -3.477390 | -1.097931 |
| **log(*M.luteus/C.afermentans*)** | 1.786337 | 0.117204 | -0.013707 | 0.098391 |
| **log(*S.aureus/C.afermentans*)** | -12.891296 | 0.000004 | -4.310889 | -2.974538 |
| **log(*S.capitis/C.afermentans*)** | 0.706425 | 0.502753 | -0.041647 | 0.077132 |
| **log(*S.epidermidis/C.afermentans*)** | 1.257191 | 0.249005 | -0.074636 | 0.244093 |
| **log(*S.hominis/C.afermentans*)** | 9.084779 | 0.000040 | 0.452786 | 0.771429 |
| **log(*S.warneri/C.afermentans*)** | 8.768258 | 0.000051 | 0.333996 | 0.580661 |
| **log(*S.mitis/C.afermentans*)** | -7.348948 | 0.000156 | -0.141701 | -0.072711 |

**Table S4.** Statistics for 24h, *in-vitro* SkinCom, RL vs Control, Related to Figure 6.

|  | **t statistics** | **p value** | **upperCI** | **lowerCI** |
| --- | --- | --- | --- | --- |
| **log(*C.acnes/C.afermentans*)** | -5.15910 | 0.00131 | -4.11286 | -1.52761 |
| **log(*M.luteus/C.afermentans*)** | 0.82765 | 0.43517 | -0.02755 | 0.05722 |
| **log(*S.aureus/C.afermentans*)** | -9.43221 | 0.00003 | -4.04021 | -2.42053 |
| **log(*S.capitis/C.afermentans*)** | -3.63578 | 0.00833 | -0.39693 | -0.08409 |
| **log(*S.epidermidis/C.afermentans*)** | 2.84383 | 0.02491 | 0.08228 | 0.89431 |
| **log(*S.hominis/C.afermentans*)** | 3.93357 | 0.00565 | 0.19773 | 0.79376 |
| **log(*S.warneri/C.afermentans*)** | 2.48534 | 0.04188 | 0.01782 | 0.71600 |
| **log(*S.mitis/C.afermentans*)** | 1.30809 | 0.23216 | -0.23427 | 0.81436 |

**Table S5.** Statistics for Day 4, SLES vs Control, Related to Figure 6.

|  | **t statistics** | **p value** | **upperCI** | **lowerCI** |
| --- | --- | --- | --- | --- |
| **log(*C.acnes/C.afermentans*)** | -1.006584 | 0.388289 | -2.011125 | 1.044617 |
| **log(*M.luteus/C.afermentans*)** | -0.247151 | 0.820740 | -2.271485 | 1.944100 |
| **log(*S.aureus/C.afermentans*)** | -0.953509 | 0.410676 | -0.886183 | 0.477579 |
| **log(*S.capitis/C.afermentans*)** | 0.229355 | 0.833340 | -2.769521 | 3.199716 |
| **log(*S.epidermidis/C.afermentans*)** | -0.075735 | 0.944398 | -2.304923 | 2.197770 |
| **log(*S.hominis/C.afermentans*)** | 0.859905 | 0.453072 | -0.345936 | 0.602096 |
| **log(*S.warneri/C.afermentans*)** | -1.075817 | 0.360825 | -1.542561 | 0.763129 |
| **log(*S.mitis/C.afermentans*)** | 0.287471 | 0.792466 | -0.252952 | 0.303188 |

**Table S6.** Statistics for Day 7, SLES vs Control, Related to Figure 6.

|  | **t statistics** | **p value** | **upperCI** | **lowerCI** |
| --- | --- | --- | --- | --- |
| **log(*C.acnes/C.afermentans*)** | -1.869453 | 0.158339 | -0.233951 | 0.060804 |
| **log(*M.luteus/C.afermentans*)** | 0.888782 | 0.439591 | -0.548265 | 0.973165 |
| **log(*S.aureus/C.afermentans*)** | 1.494316 | 0.231953 | -0.177493 | 0.491722 |
| **log(*S.capitis/C.afermentans*)** | 1.762337 | 0.176220 | -0.475287 | 1.654939 |
| **log(*S.epidermidis/C.afermentans*)** | 3.206298 | 0.049095 | 0.007707 | 2.064444 |
| **log(*S.hominis/C.afermentans*)** | 2.089460 | 0.127843 | -0.170947 | 0.824543 |
| **log(*S.warneri/C.afermentans*)** | 1.337519 | 0.273415 | -0.592327 | 1.451166 |
| **log(*S.mitis/C.afermentans*)** | -0.535378 | 0.629533 | -0.216667 | 0.154266 |

**Table S7.** Statistics for Day 4, RL vs Control, Related to Figure 6.

|  | **t statistics** | **p value** | **upperCI** | **lowerCI** |
| --- | --- | --- | --- | --- |
| **log(*C.acnes/C.afermentans*)** | -0.453956 | 0.665819 | -0.912310 | 0.626776 |
| **log(*M.luteus/C.afermentans*)** | 0.876440 | 0.414499 | -0.365185 | 0.772787 |
| **log(*S.aureus/C.afermentans*)** | -0.273301 | 0.793786 | -0.603805 | 0.482476 |
| **log(*S.capitis/C.afermentans*)** | 0.715330 | 0.501279 | -0.464750 | 0.848735 |
| **log(*S.epidermidis/C.afermentans*)** | 0.364150 | 0.728235 | -1.049873 | 1.416992 |
| **log(*S.hominis/C.afermentans*)** | 0.019290 | 0.985235 | -0.766951 | 0.779140 |
| **log(*S.warneri/C.afermentans*)** | -0.142990 | 0.890980 | -0.683197 | 0.607758 |
| **log(*S.mitis/C.afermentans*)** | 0.258699 | 0.804514 | -0.785381 | 0.971083 |

**Table S8.**
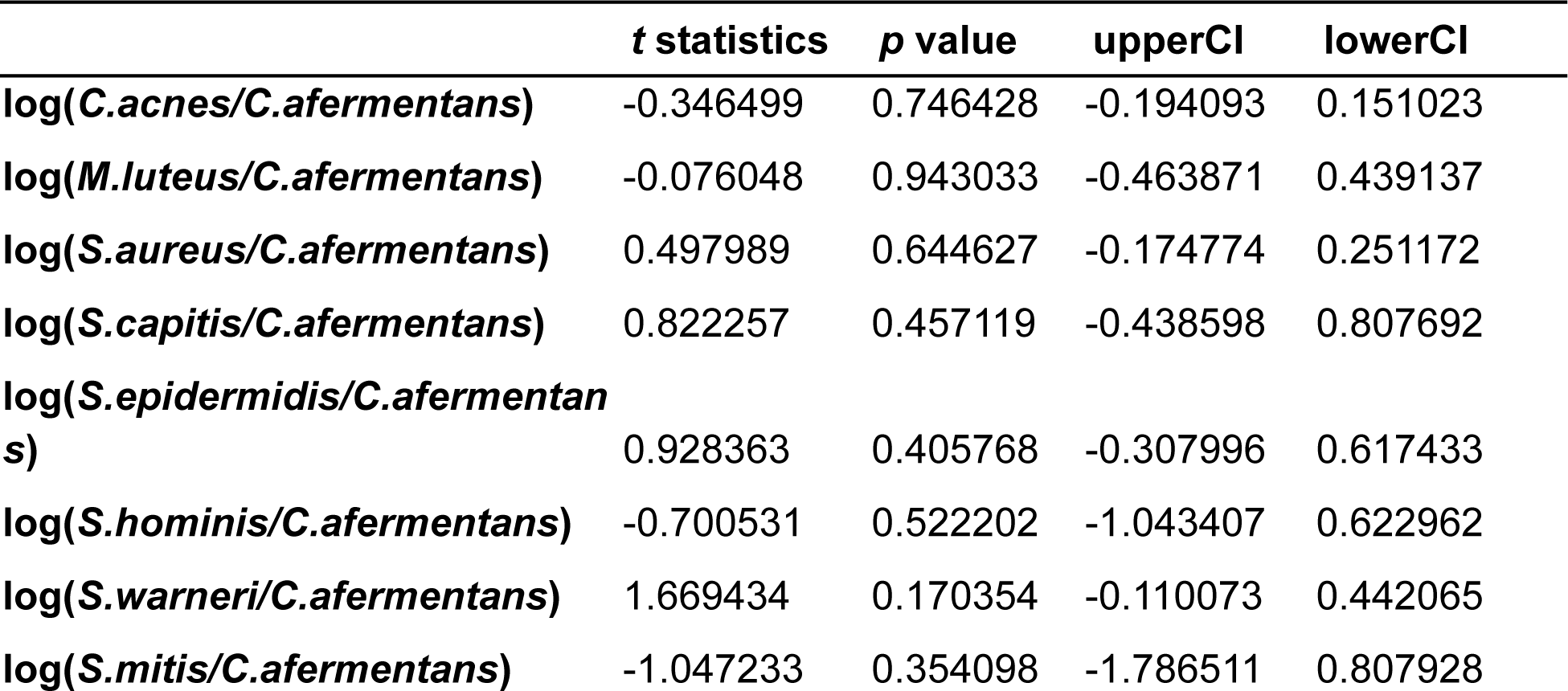
Statistics for Day 7, RL vs Control, Related to Figure 6.

